# Superoxide enters neurons via LRRC8A – containing volume-regulated anion channels

**DOI:** 10.1101/2024.02.20.580732

**Authors:** Kate Harris, Gokhan Uruk, Seok Joon Won, Nguyen Mai, Paul Baxter, Katharina Everaerts, Rajan Sah, Raymond A. Swanson

**Affiliations:** Dept of Neurology University of California, San Francisco; Neurology Service, San Francisco Veterans Affairs health System; Institute of Neurobiology, Heinrich Heine University Düsseldorf, Düsseldorf, Germany; Department of Internal Medicine, Cardiovascular Division, Washington University School of Medicine, St. Louis, MO, USA; John Cochran VA Medical Center, St. Louis, MO, USA

## Abstract

Superoxide (O_2_**^-^**) is both an intercellular signaling molecule and a cause of neuronal oxidative stress. Superoxide entry into neurons is thought to be indirect, requiring its dismutation to nonpolar hydrogen peroxide. Here we show instead that superoxide enters neurons directly, via LRRC8A-containing volume-sensitive organic anion channels. In primary cultures, neuronal oxidative stress induced either by NMDA receptor stimulation or exposure to authentic superoxide was blocked by the anion channel blockers DIDS and DCPIB and by LRRC8A gene disruption. In mouse cortex, neuronal oxidative stress induced by either NMDA injection or transient ischemia was likewise blocked by both DCPIB and LRRC8A gene disruption. These findings identify a role for LRRC8A-containing volume-sensitive organic anion channels in neuronal oxidative signaling, stress, and glutamate excitotoxicity.

## INTRODUCTION

The anionic free radical superoxide (O_2_^-^) serves as an intercellular signaling molecule in the central nervous system (Aizenman et al., 1990; De Pasquale et al., 2014; Klann et al., 1998; Terzi and Suter, 2020) and can also contribute to neuronal injury. Although not itself highly reactive, superoxide combines with other molecules to form extremely reactive species such as peroxynitrite and hydroxyl radical (Beckman and Koppenol, 1996). High rates of superoxide production, as occur during neuronal excitotoxicity, can thereby lead to neuronal oxidative stress and death (Brennan et al., 2009; Lafon-Cazal et al., 1993; Patel et al., 1996; Reyes et al., 2012). Superoxide is produced both intracellularly, as a by-product of mitochondrial oxidative metabolism and other processes (Hamanaka and Chandel, 2010), and extracellularly by members of the NADPH oxidase (NOX) family of enzymes (Sorce and Krause, 2009). Neurons express NADPH oxidase-2 (NOX2) at their plasma membranes (Keeney et al., 2022; Tejada-Simon et al., 2005), with neuronal NOX2 stimulated by NMDA (N-methyl-d-aspartate)-type glutamate receptor activation, prion protein binding, and other factors (Adams et al., 1989; Brennan-Minnella et al., 2013; Brennan et al., 2009; Dugan et al., 2009; Schneider et al., 2003; Walsh et al., 2014). Superoxide produced by NOX2 at the cell membrane is released into the extracellular space, but how extracellular superoxide gains entry into neurons has been unresolved.

As a charged molecule, superoxide cannot readily cross cell membranes. Many authors assume that it must first be converted spontaneously or by action of superoxide dismutase to non-polar hydrogen peroxide (H_2_O_2_), which can then cross lipid membranes passively or via aquaporin channels (Bienert et al., 2007). However, this view is inconsistent with observations that extracellular superoxide dismutase reduces, rather than increases, the oxidative stress and cell injury induced by superoxide (Buczynski et al., 2018; Reyes et al., 2012; Sun et al., 2019).

Anion channels have been identified as a route of superoxide influx in certain non-neuronal cell types, including erythrocytes (Lynch and Fridovich, 1978), amniotic cells (Ikebuchi et al., 1991) and endothelial cells (Hawkins et al., 2007). Neurons express a variety of anion channels, including volume-regulated anion channels (VRACs). VRACs are so-named because they were first identified by their role in the cellular response to hypo-osmolar cell swelling, in which they permit efflux of Cl^-^ and other anions followed by water (Chen et al., 2019), but under physiological conditions VRACs are activated by other stimuli (Chen et al., 2019; Pedersen et al., 2015) and function in a variety of cell processes (Kumar et al., 2020; Kang et al., 2018; Alghanem et al., 2021; Gunasekar et al., 2023). VRACs are formed as heteromers of LRRC8 proteins (LRRC8A-E), of which LRRC8A is essential for channel function (Qiu et al., 2014; Voss et al., 2014). Different LRRC8 subunit compositions yields different channel properties and substrate selectivity (Schober et al., 2017), and in various cell types VRACs have been found to nonspecifically conduct anions including chloride, taurine, and glutamate (Osei-Owusu et al., 2018; Yang et al., 2019).

VRAC inhibitors have long been recognized as neuroprotective in rodent stoke models (Balkaya et al., 2023; Feng et al., 2004; Kimelberg et al., 2000; Zhang et al., 2008). This effect was initially ascribed to inhibition of excitotoxic glutamate release from astrocyte VRACs (Phillis et al., 1998; Seki et al., 1999; Zhang et al., 2008); however, VRACs are a relatively minor route of glutamate efflux during ischemia (Rossi et al., 2000; Swanson et al., 2004), and more recent work shows a poor correlation between VRAC-mediated glutamate release and ischemic neuronal injury (Balkaya et al., 2023).

Here we show that LRRC8A-containing VRACs gate entry of superoxide into neurons. This observation provides an alternative explanation for how VRACs are involved in ischemic and excitotoxic neuronal injury, and more broadly suggests that the location and functional state of VRACs may be a critical factor determining both neuronal superoxide signaling and superoxide-mediated injury.

## MATERIALS AND METHODS

### Animals

Studies were approved by the Institutional Animal Care and Use Committee at the San Francisco Veterans Affairs Medical Center and were performed in accordance with the National Institutes of Health Guide for the Care and Use of Laboratory Animals. Results are reported in accordance with the ARRIVE guidelines. C57Bl/6 mice were obtained from Jackson Laboratories. Breeding pairs of LRRC8A floxed/floxed (LRRC8A^fl/fl^) mice on the C57BL/6 background (Zhang et al., 2017) were a gift from Rajan Sah, Washington University, St Louis. Breeding pairs of Clcn3^+/-^ mice (Arreola et al., 2002) were a gift from Fred Lamb, Vanderbilt University. Studies using adult mice used equal numbers of both sexes, aged 3-6 months.

### Cell Culture Studies Neuron cultures

Primary cultures of cortical neurons were obtained from embryonic day 16 C57BL/6 wild-type, CLCn3^+/-^, or LRRC8A^fl/fl^ mice as described previously (Ghosh et al., 2021). Neurons were plated on 24 well plates on poly-D-lysine–coated glass coverslips and maintained in serum-free NeuroBasal medium (Gibco) containing 5 mM glucose at 37° C in 5% CO_2_. The CLCn3^-/-^ embryos were generated by crossing CLC3^+/-^ mice, genotyped after culture preparation.

### LRRC8A downregulation

Cultured LRRC8A^fl/fl^ neurons were transduced with a Cre-encoding lentivirus (LV-synapsin(Syn)-Cre-GFP; SignaGen) at a multiplicity of infection (MOI) of 20 to suppress LRRC8A expression. A virus encoding GFP but not Cre was used as a control (LV-Syn-GFP). LRRC8A gene expression was assessed 7 days after transduction by RNAscope *in situ* hybridization (Advanced Cell Diagnostics) using the Mm-Lrrc8a probe (Cat# 458371). Under the conditions employed, the PPIB positive control (Cat# 320881) gave a signal in every cell, and the dapB negative control (Cat# 320871) gave no signal. The number of LRRC8A RNAscope dots was counted in GFP-positive neurons to assess efficacy of LRRC8A downregulation. LRRC8A protein expression was assessed 7 days after transfection by western blotting, performed as described (Ghosh et al., 2021). The blots were probed with rabbit polyclonal anti-LRRC8A antibody (1:1000; Bethyl Labs, #A304-175A) overnight at 4 °C followed by incubation with HRP-conjugated secondary antibodies (1:5000; Cytiva Life Sciences, NA934V/NA931V). Mouse monoclonal anti-β-actin (1:7500, ThermoFisher, MA1-140) was used as a loading control. Blots were visualized using SuperSignal™ West Pico PLUS chemiluminescent substrate, and the integrated band optical densities were analyzed using ImageJ.

### KO_2_ ^-^ and NMDA exposures

Experiments were begun by replacing the culture medium with a balanced salt solution (BSS) containing 1.2 mM CaCl_2_, 0.8 mM MgSO_4_, 5.3 mM KCl, 0.4 mM KH_2_PO_4_, 137 mM NaCl_2_, 0.3 mM NaHPO_4_, 5 mM glucose and 10 mM 1,4-piperazinediethanesulfonate buffer (final pH 7.25). The MgSO_4_ concentration was reduced to 0.4 mM for experiments conducted with NMDA. KO_2_^-^ studies were performed with neurons on day 7-8 *in vitro* and NMDA studies were performed on day 13-14, to allow time for the development of NR2B-containing NMDA receptors (Zhong et al., 1994).

KO_2_ ^-^ was prepared as a saturated solution in DMSO and added to the BSS as 10 µL DMSO / 400 µl medium (Fig. 1A) to produce an initial O_2_^-^ concentration of approximately 40 µM (Hawkins et al., 2007; Reiter et al., 2000; Reyes et al., 2012). NMDA was prepared in water and used at a final concentration of 100 µM. Where indicated, the following chemicals were added to the culture medium 10 minutes prior to the addition of KO_2_^-^ or NMDA; diisothiocyanatostilbene-2,2′-disulfonic acid (DIDS; MilliporeSigma), 4-(2-Butyl-6,7-dichloro-2-cyclopentyl-indan-1-on-5-yl) oxobutyric acid (DCPIB; Tocris), superoxide dismutase (SOD; MilliporeSigma), catalase (MilliporeSigma) or GSK2795039 (Cayman Chemical). Dihydroethidium (DHE; Invitrogen) was added at 2 µM final concentration immediately prior to KO_2_ ^-^ or NMDA. Following 20-minute incubations with KO_2_ ^-^ or NMDA, the cells were rinsed in phosphate-buffered saline and fixed in 4% paraformaldehyde solution (PFA) for 5 minutes.

**Figure 1.**
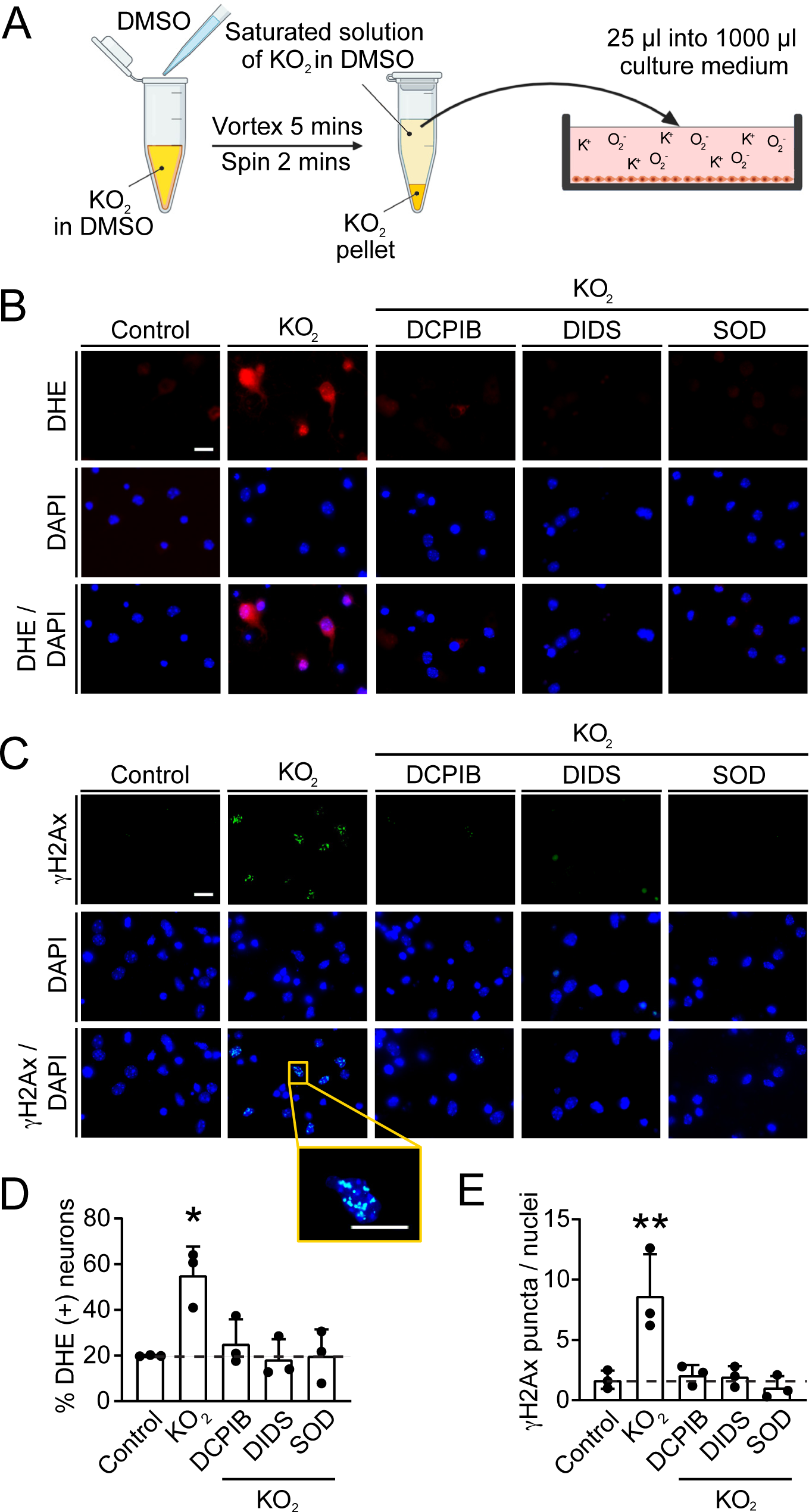
Anion channel blockers prevent superoxide entry into cultured neurons. **A.** Delivery of superoxide (O_2_^-^) to culture medium using saturated KO_2_^-^ in DMSO. **B.** Oxidized DHE (red) in primary cortical neurons after incubation with KO_2_^-^. Cell nuclei are labeled with DAPI (blue). Where indicated, cultures were co-incubated with superoxide dismutase (SOD, 1 U/ml), DCPIB (10 μM), or DIDS (100 μM). **C.** Foci of γH2Ax formation (green) in neurons treated as in (A). Inset shows high magnification view of nuclei with γH2Ax puncta. Scale bars = 10 µm. **D,E.** Quantification of oxidized DHE and γH2Ax signals. *p < 0.05, **p < 0.01 versus KO_2_^-^ alone by Dunnett’s test (n = 3).

### Immunostaining and superoxide detection

Immunostaining was performed as described in (Ghosh et al., 2021), using rabbit anti-phospho-Histone H2A.X Ser139 (1:500, Cell Signaling, #2577). The coverslips were mounted with DAPI to counterstain cell nuclei. Photography and image analysis was done by individuals blinded to the experimental conditions. Photomicrographs were prepared from 3 arbitrary regions of comparable neuronal density in each well. The DHE and γH2Ax signals were analyzed in neuronal soma as defined by DAPI staining or, in lentivirus-transduced neurons, by GFP. The mean integrated density of DHE and γH2Ax was measured in each neuron and expressed as percent of neurons in which the signal exceeded the 80th percentile signal of neurons in the control wells of the same 24-well plate (Brennan et al., 2009). For some experiments, the number of γH2Ax foci per nuclei was calculated in place of mean integrated fluorescence because the small size of individual puncta did not produce significant increases in overall fluorescence intensity.

### In vivo methods

#### Virus and drug administration

Studies used C57BL/6 wild-type or LRRC8A^fl/fl^ mice, age 3.5 - 5 months, with equal numbers of male and female mice allotted to each experimental condition. Stereotaxic injections of virus were made AP +1.0, ML −2.0, DV −1.0 mm from Bregma under isoflurane anesthesia with AAV9-Syn-Cre-Syn-GFP (SignaGen, #SL116013) at a titer of 1x 10^12^ / ml in 0.5 µl saline. Bupivacaine and buprenorphine SR were injected subcutaneously and intraperitoneally to provide pain relief. The virus-injected mice were used for experiments 4 weeks after the injections.

Intracortical NMDA injections (0.6 μl of 10 mM NMDA in saline) were made to the same coordinates used for the virus injections, with control mice receiving saline vehicle only (Fig. 6a). DCPIB was administered intracerebroventricularly (i.c.v.; AP −0.2, ML −1.0, DV −2.25 mm) as 2 µl of 5 mM DCPIB in 10% DMSO / saline (Taylor et al., 2021; Zhang et al., 2008), with control mice receiving 10% DMSO/saline only. The DCPIB injections were completed 20 minutes prior to intracortical NMDA or saline injections. DHE was administered intraperitoneally 10 mg/kg in 10% DMSO / saline 15 minutes prior to intracortical NMDA or saline injections. For tMCAo, DHE was administered 5 minutes prior to occlusion and 1, 2, and 3 hours after reperfusion (Fig 7A).

### Transient middle cerebral artery occlusion (tMCAo)

Adult (age 3–6 months) male and female mice were subjected to tMCAo using the Weinstein intraluminal filament method (Longa et al., 1989) under isoflurane anesthesia as described (Won et al., 2018). Cortical blood flow was monitored using laser Doppler flowmetry (Moor instruments Ltd) to confirm ischemia. Blood flow was restored by retracting the monofilament after 45 minutes. Mice were excluded from analysis if blood flow did not drop below 25% of baseline or return above 80% of baseline. Mice were euthanized for brain harvest 3 hours and 15 minutes after reperfusion.

### Histology and image analysis

Anesthetized mice were cardio-perfused with saline followed by buffered 4% paraformaldehyde solution (PFA) and the brains were cryostat sectioned. Antigen retrieval was performed by incubating sections in 10 mM sodium citrate, pH 6.0, at 80 °C for 30 minutes. Immunostaining was performed using rabbit anti-phospho-Histone H2A.X Ser139 (1:500; Cell Signaling, #2577), mouse anti-NeuN (1:500; MilliporeSigma #MAB377B) and SOX99 (1:500; MilliporeSigma #MAB5535).

Fluorescent secondary antibodies were obtained from Life Technologies and used at a 1:500 dilution. Photographs were taken by confocal microscopy from 4 predetermined locations on each of two consecutive coronal sections of each brain. For the NMDA-injected brains the photographs were taken 250 µm from the needle track, and for the tMCAo brains they were taken 250 µm from the infarct border (as identified by NeuN immunoreactivity). DHE fluorescence and γH2Ax foci were quantified in neuronal soma using ImageJ. The soma were identified by DAPI or, where virus transduction was done, by GFP. The mean integrated densities of DHE and γH2Ax fluorescence were measured in each neuron and expressed as the percent of neurons in which the signal intensity exceeded the 80^th^ percentile value of neurons measured in control brains processed in parallel.

### RNAscope *in situ* hybridization

RNAscope assays were performed as described above for the cell culture studies, using 4 µm optical sections captured by spinning disc confocal microscopy. The number of RNAscope dots was counted in each neuron (as defined by NeuN labeling) using ImageJ, and the mean intensity of the GFP signal was measured on the same NeuN-defined regions using ImageJ.

### Statistics

The “n” of each study was defined as the number of mice or, for cell cultures, the number of independent experiments. Each independent experiment contained triplicate or quadruplicate culture wells. GraphPad Prism was used for statistical analysis and to generate graphs. Numerical data were expressed as means ± SEM and analyzed using one-way ANOVA and either the Tukey–Kramer test where multiple groups were compared against one another, or Dunnett’s test where multiple groups were compared against a common control group. Where only two groups were compared, the two-sided t-test was used. P values < 0.05 were considered statistically significant.

## RESULTS

### Extracellular superoxide enters neurons via volume-regulated anion channels (VRACs)

We first used pharmacological approaches to confirm that extracellular superoxide enters neurons via anion channels. For these studies, primary neuronal cultures were exposed to O_2_^-^ delivered as a saturated solution of KO_2_^-^ in DMSO (Fig. 1A). KO_2_^-^ dissociates to K^+^ and O_2_^-^ when added to the aqueous culture medium, providing an initial O_2_^-^ concentration of approximately 40 µM (Reiter et al., 2000). Intracellular superoxide was measured using the fluorogenic probe dihydroethidium (DHE), which upon oxidation binds to DNA and generates red fluorescence (Chen et al., 2013). Neurons incubated with KO_2_^-^ exhibited a large increased in intracellular DHE fluorescence, and this increase was suppressed by co-incubation with either the non-specific anion channel blocker DIDS (Friard et al., 2017) or the LRRC8A channel inhibitor DCPIB (Decher et al., 2001) (Fig. 1). Crucially, the DHE signal induced by KO_2_^-^ was also suppressed by co-incubation with superoxide dismutase (SOD), thus confirming that the DHE signal was attributable to superoxide entering cells rather than from H_2_O_2_ formed from superoxide in the culture medium.

As a marker of superoxide-induced oxidative stress in neurons we assessed formation of γH2Ax foci, which accumulate at sites of double stranded DNA breaks (Mah et al., 2010). Results of these studies paralleled those using DHE; the formation of γH2Ax foci was inhibited by SOD, DIDS, and DCPIB (Fig. 1B). Additional studies confirmed that catalase, which degrades H_2_O_2_, did not eliminate the KO_2_^-^ induced DHE and γH2Ax signals (Suppl. Fig. 1). Moreover, neither DIDS nor DCPIB blocked the signals induced by H_2_O_2_ (Suppl. Fig. 2). These findings further excluded the possibility that the DHE and γH2Ax signals were produced by H_2_O_2_ formed in the medium. Given that glutamate can exit cells via VRACs, we also considered the possibility that KO_2_^-^ was indirectly causing egress of glutamate via neuronal VRACs, and that the observed effects of that anion channel inhibitors and SOD were attributable to preventing this release. However, this possibility was excluded by the absence of DHE oxidation or γH2AX formation in cultures (also D.I.V. 7) to which 100 µM glutamate was added (Suppl. Fig. 3). Together these results establish that extracellular superoxide enters neurons via VRACs.

We next evaluated a physiological source of superoxide, neuronal NOX2, which releases superoxide to the extracellular space in response to NMDA receptor activation (Brennan-Minnella et al., 2013; Brennan et al., 2009; Reyes et al., 2012). Neuronal cultures treated with NMDA showed an increase in DHE fluorescence, and this was blocked by the NOX2 inhibitor GSK2795039 (Suppl. Fig. 4). As observed using KO_2_ ^-^ as the source of superoxide, the increased DHE fluorescence and γH2Ax foci formation induced by NMDA was attenuated by both SOD and the anion channel blockers DIDS and DCPIB (Fig. 2).

**Figure 2.**
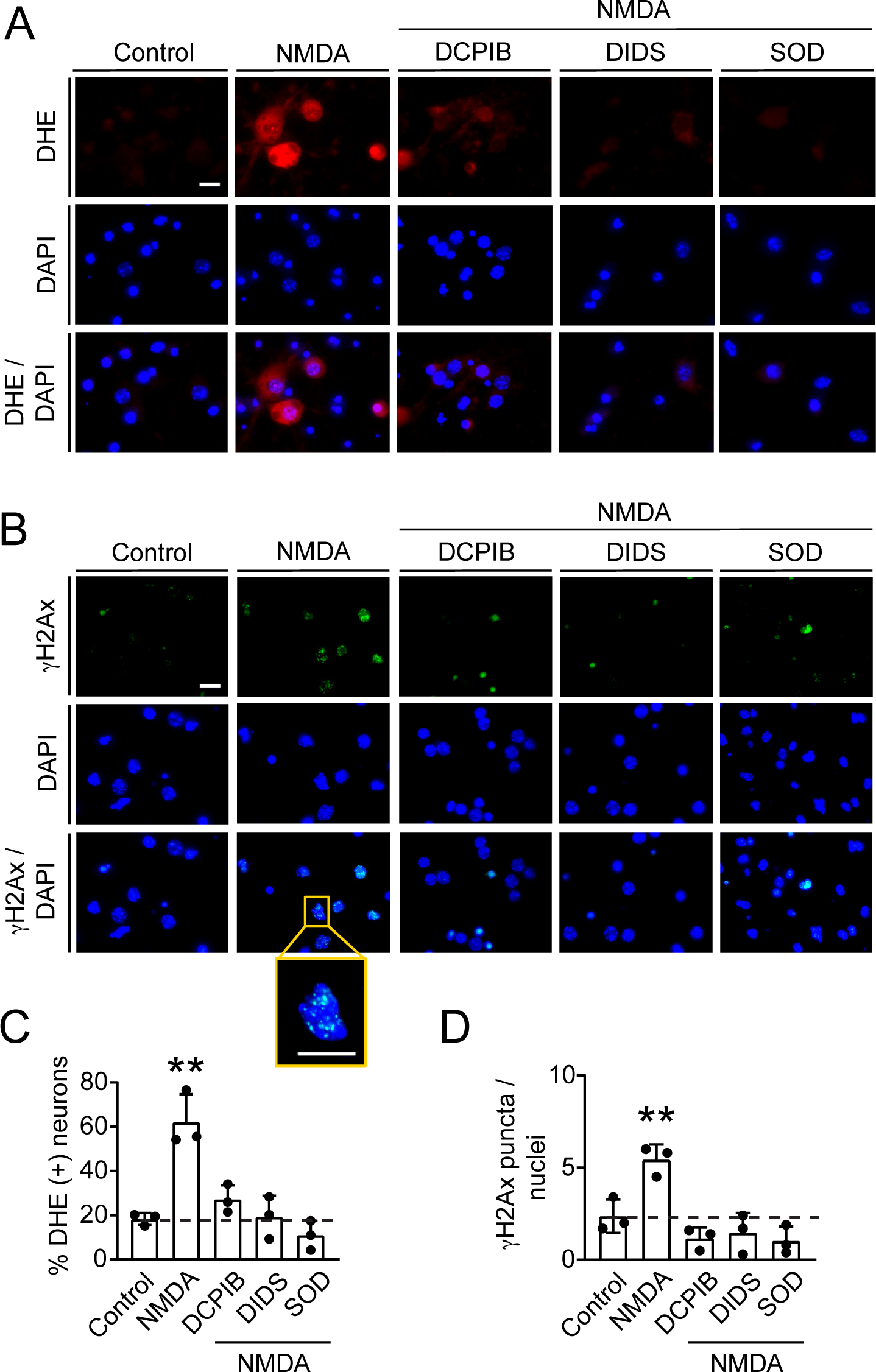
Anion channel blockers prevent NMDA-induced oxidative stress in cultured neurons. **A.** Oxidized DHE (red) in primary cortical neurons treated with NMDA. Cell nuclei are labeled with DAPI (blue). Cultures were co-incubated with superoxide dismutase (SOD; 1 U/ml), DCPIB (10 μM), or DIDS (100 μM) as indicated. **B.** Foci of γH2Ax formation (green) in neurons treated as in (A). Inset shows high magnification view of nuclei with γH2Ax puncta. **C,D)** Quantification of oxidized DHE and γH2Ax. **p < 0.01 versus NMDA alone by Dunnett’s test (*n* = 3).

### VRAC gene disruption prevents superoxide transit into primary neurons

To confirm VRACs as the channels mediating superoxide entry, we generated neurons in which the essential VRAC subunit LRRC8A, was deleted. Neuronal cultures from LRRC8A^fl/fl^ mice were incubated with lentirvirus encoding both Cre recombinase and GFP (LV-Cre-GFP). Resultant knockdown of LRRC8A expression was confirmed using RNAscope *in situ* hybridization and western blots (Fig. 3A-D). The LRRC8A^fl/fl^ neurons treated with LV-Cre-GFP showed no increase in DHE signal after KO_2_^-^ treatment, whereas LRCC8A^fl/fl^ neurons treated with the control lentivirus encoding GFP-only (no Cre) showed the expected large increase in DHE oxidation after exposure to KO_2_^-^ (Fig. 3). Studies using γH2Ax formation as a marker of intracellular oxidative stress showed a similar result, with γH2Ax formation likewise attenuated in the LRCC8A^fl/fl^ neurons transfected with LV-Cre-GFP (Fig. 3). We then investigated the effect of LRRC8A deletion on NMDA - induced superoxide entry and oxidative stress. In parallel to the results obtained using DHE as a marker of oxidative stress, LRCC8A^fl/fl^ neurons treated with LV-Cre-GFP showed both reduced DHE signal and reduced γH2Ax foci formation compared to that observed in the LV-GFP – treated controls (Fig. 4).

**Figure 3.**
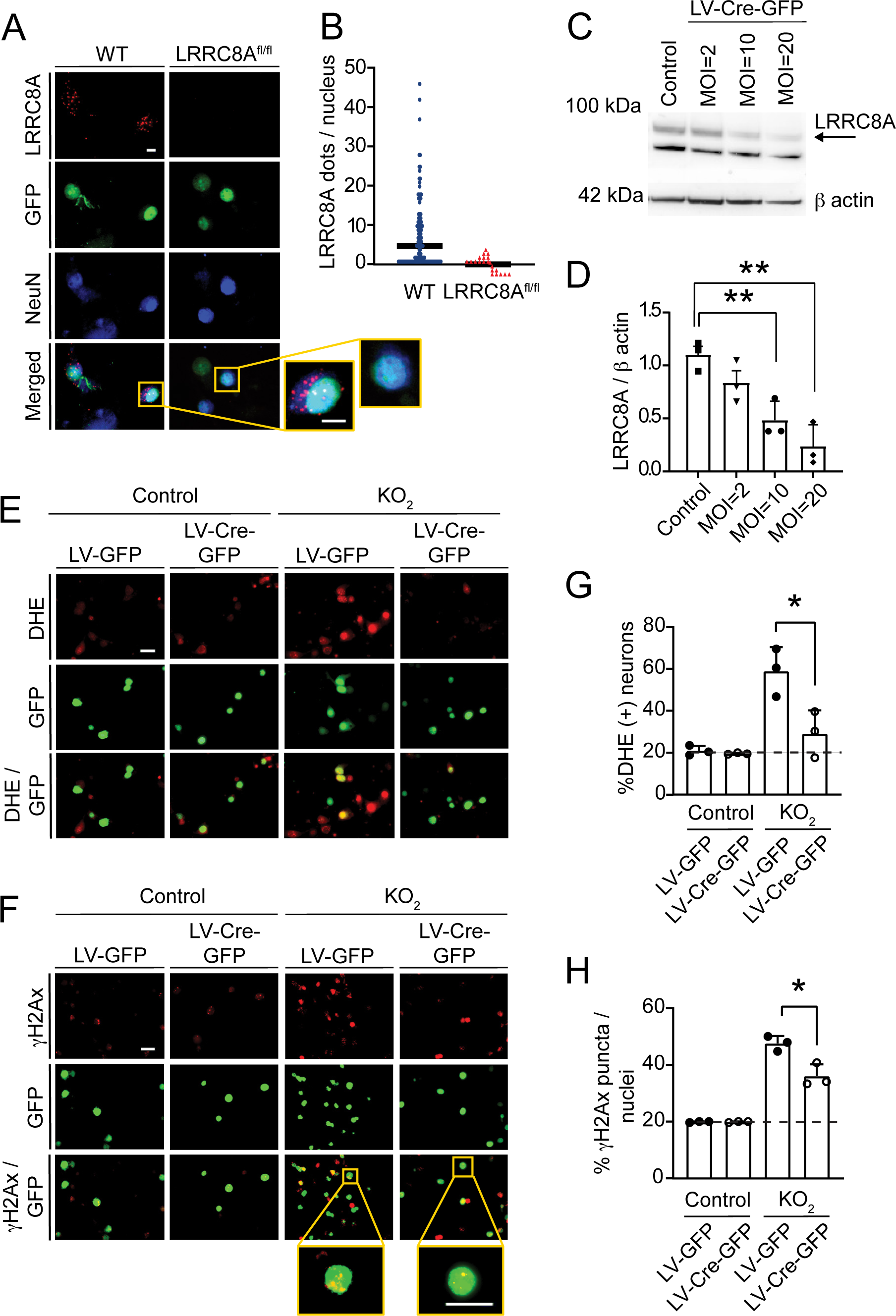
LRRC8A deletion prevents superoxide - induced oxidative stress in cultured neurons. **A.** RNAscope images of primary neurons from LRRC8A^fl/fl^ and WT mice transduced with LV-synapsin-Cre-GFP (MOI 20) and stained for LRRC8A mRNA (red). Insets show higher magnification. Dots represent values from individual neurons, n=2 mice for each genotype. **B.** Quantification of LRRC8A mRNA shown in (A). **C.** Western blot showing LRRC8A in LRRC8A^fl/fl^ neuronal cultures exposed to LV-synapsin-Cre-GFP. **D.** Quantification of western blots shown in (C). **E.** Oxidized ethidium (DHE, red) in primary neurons transfected with LV-GFP (control) or LV-GFP-Cre and exposed to KO_2_^-^ five days later. GFP expression (green) shows virus transduction. **F.** γH2Ax foci (red) in neurons treated as in (E). Inset shows high magnification view of nuclei with γH2Ax puncta. **G,H.** Quantification of oxidized DHE and γH2Ax puncta. *p < 0.05 versus LV-GFP by Student’s t-test (*n* = 3).

**Figure 4.**
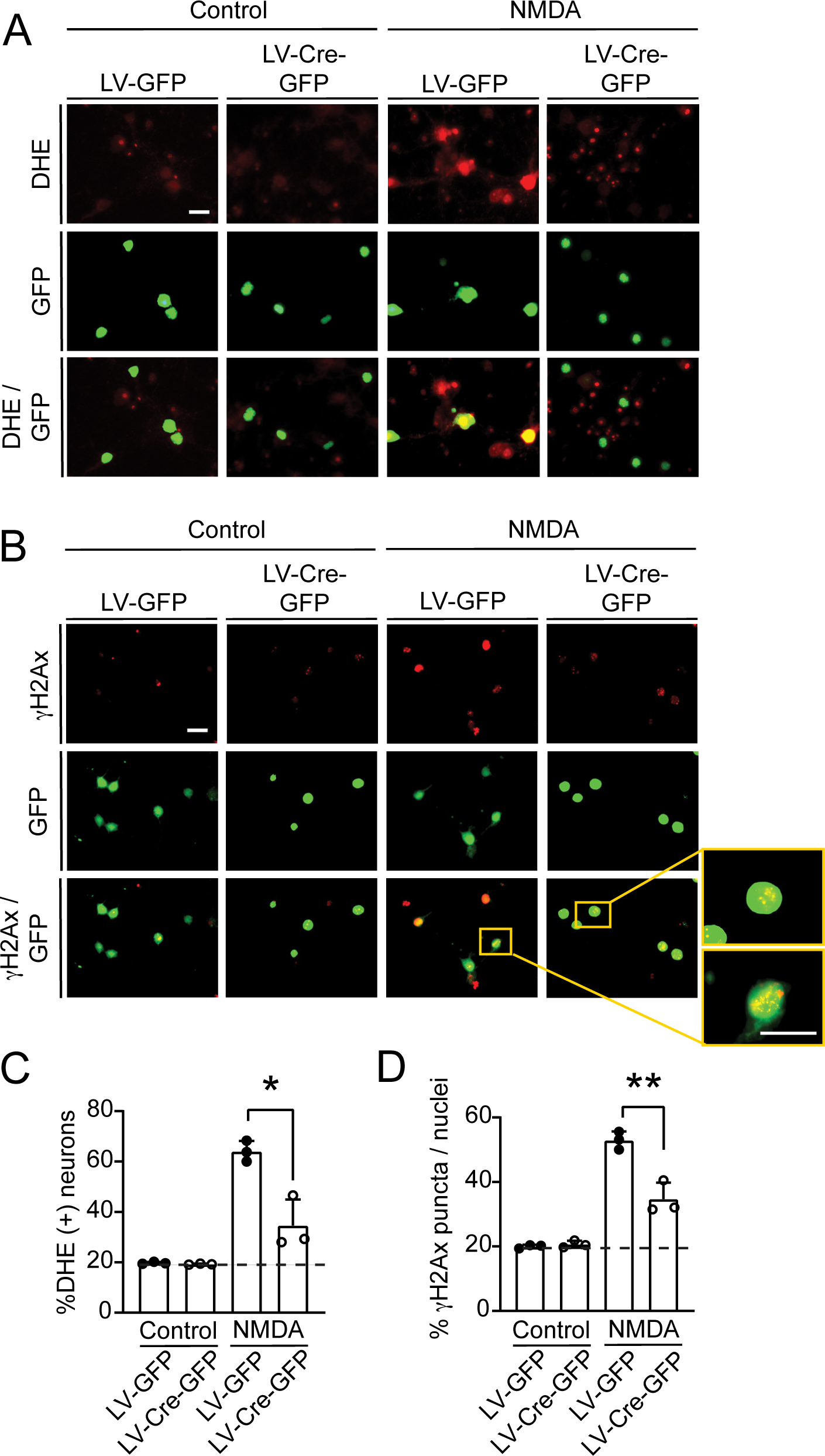
LRRC8A deletion prevents NMDA-induced oxidative stress in cultured neurons. **A.** Oxidized DHE (red) in LV-GFP (control) or LV-GFP-Cre transfected neurons following incubation with NMDA. **B.** Images of γH2Ax foci (green) in neurons treated as in (A). Inset shows high magnification view of nuclei with γH2Ax puncta. **C,D.** Quantification of oxidized DHE and γH2Ax. **p < 0.01 and *p < 0.05 versus LV-GFP by Student’s t-test (*n* = 3).

With both KO_2_^-^ and NMDA, the effects of LRRC8A gene deletion on superoxide entry and oxidative stress were in some cases less complete than observed with the anion channel inhibitors, suggesting the possibility that other channels besides VRACs may contribute to the superoxide conductance. Given that the ClC-3 chloride channel has been identified as a superoxide-conducting channel in endothelial cells (Hawkins et al., 2007), we evaluated the possibility that ClC-3 might also gate superoxide influx into neurons. However, neurons cultured from mice deficient in CLC-3 (*ClCn3*^-/-^ mice) showed no decrement in either KO_2_ ^-^ or NMDA - induced DHE oxidation (Suppl. Fig. 4).

### Requisite role for neuronal VRACs in NMDA- and ischemia-induced neuronal oxidative stress in vivo

We next evaluated the role of LRRC8A - containing VRACs in superoxide entry into neurons in mature brain, using NMDA injections into mouse parietal cortex. The NMDA injections produced a large neuronal DHE signal, as expected, and this signal was completely suppressed by prior intracerebroventricular DCPIB injection (Fig. 5). The NMDA-injected cortices also showed a large increase in γH2Ax formation, which was likewise suppressed by DCPIB (Fig. 5).

**Figure 5.**
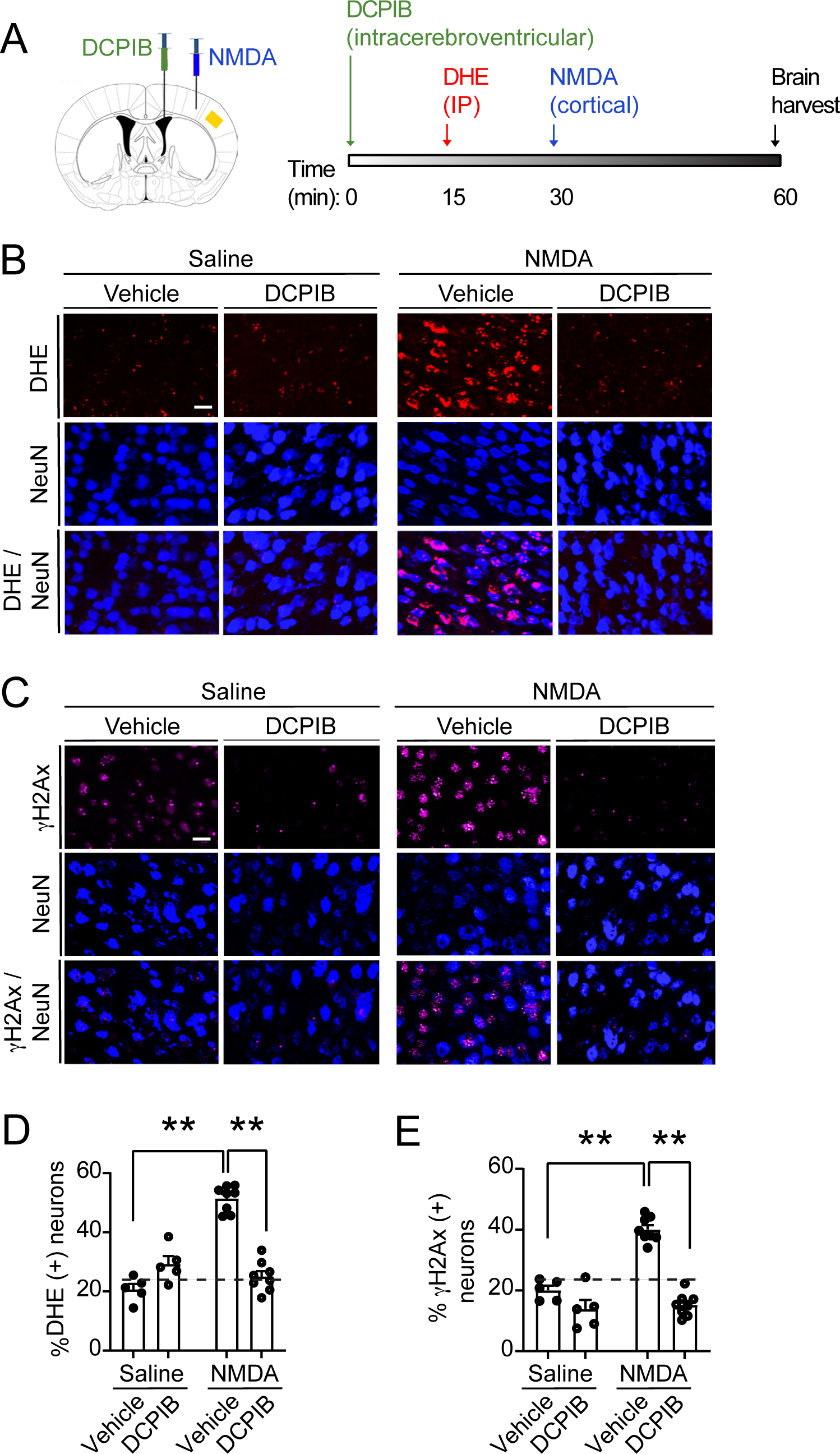
DCPIB prevents oxidative stress after intracortical NMDA injection. **A.** Time course of DCPIB, DHE, and NMDA injections. Red rectangle depicts the photographed region. **B,C.** Oxidized DHE (red) and γH2Ax foci (magenta) in neurons (NeuN, blue) following intracortical NMDA injections. Controls received intracortical vehicle (water) injections. Where indicated, mice were pre-treated with intacerebroventricular injection of DCPIB or 10% DMS vehicle. Scale bar = 20 µM. **D,E.** Quantification of oxidized DHE and γH2Ax foci in the neurons. n = 5-8; *p < 0.05, **p < 0.01 by one-way ANOVA with Tukey’s test.

We then evaluated the effect of VRAC gene disruption on the NMDA-induced neuronal oxidative stress in mouse brain cortex. Genetic disruption of VRACs was accomplished using AAV9-Synapsin(Syn)-Cre-Syn-GFP injection into cortices of LRRC8A^fl/fl^ mice with wild-type mice injected with the same virus serving as controls. RNAscope analysis performed 4 weeks after virus injection showed a near-complete suppression of LRRC8A gene expression in the transfected (GFP-expressing) neurons of LRRC8A^fl/fl^ mice and confirmed that astrocytes were not transfected (Fig. 6A-D). NMDA injected into cortex of the LRRC8A^fl/fl^ mice in which neuronal LRRC8A had been disrupted by AAV-Syn-Cre-Syn, showed markedly reduced DHE signal relative to wild-type mice treated identically (Fig. 6F, G). The DNA damage marker γH2Ax likewise showed reduced signal in the LRRC8A gene-disrupted neurons (Fig. 6H, I).

**Figure 6.**
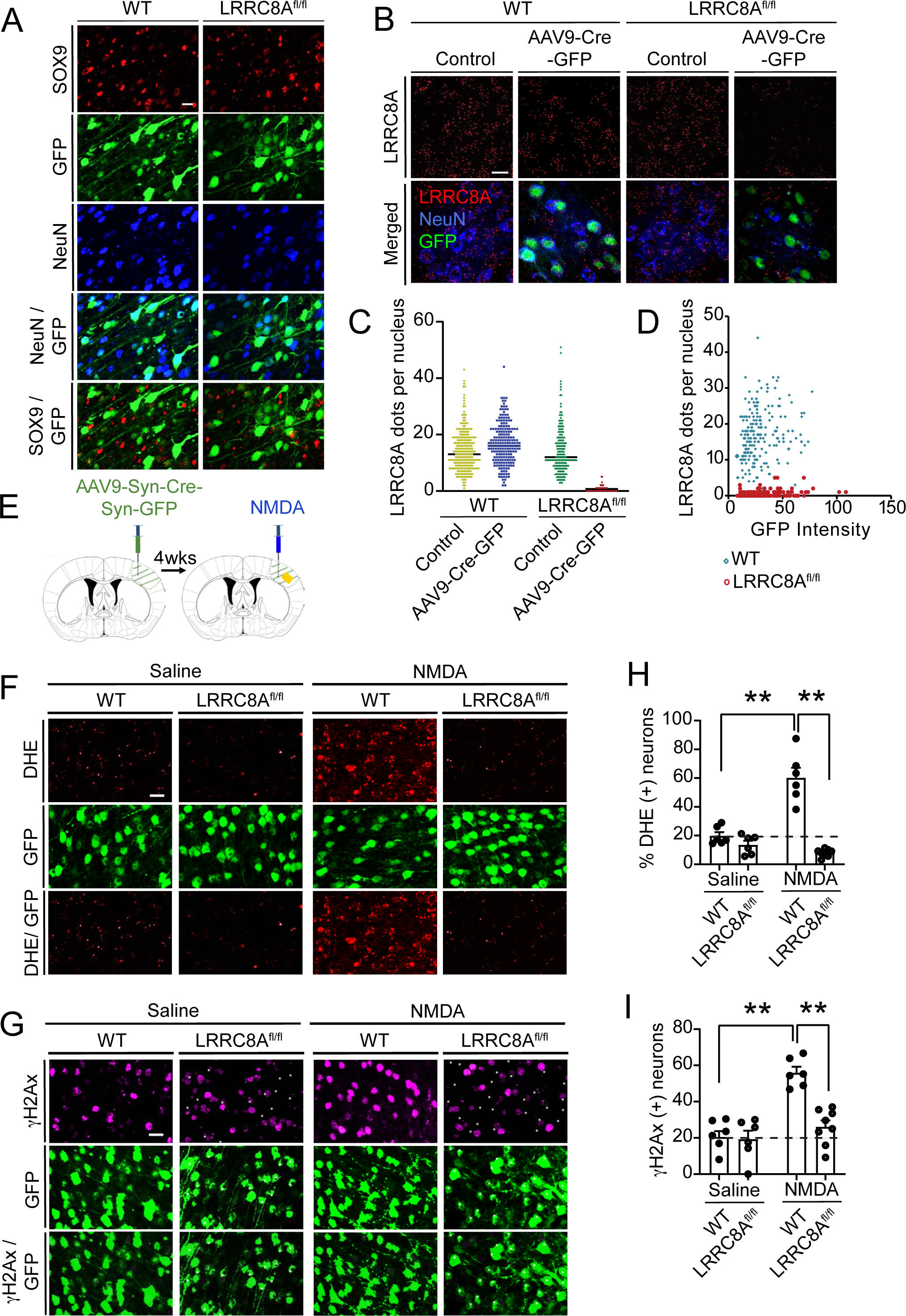
LRRC8A deletion prevents oxidative stress after NMDA injection. **A.** AAV9 injected cortex showing co-localization of the GFP signal with the neuronal marker NeuN but not the astrocyte marker Sox9 in both WT and LRRC8A^fl/fl^ mice. **B**. RNAscope visualization of LRCC8A mRNA shows Cre-lox induced gene disruption in transduced LRRC8A^fl/fl^ neurons. C. Quantification of LRRC8A mRNA. D. Plot of LRRC8A signal vs. GFP signal intensity shows near-absence of LRRC8A signal in cells with any detectable GFP signal. Dots represent values from individual neurons, n=2 mice for each genotype. **E.** Diagram of AAV and NMDA injections. Red rectangle depicts the photographed region). **F,G.** Representative images of oxidized DHE (red) and γH2Ax foci (magenta) in AAV9-Syn-GFP-Syn-cre transfected neurons of WT or LRRC8A^fl/fl^ mice following intracortical NMDA injections. Scale bar = 20 µM. H,I. Quantification of oxidized DHE and γH2Ax in the GFP(+) neurons. **E.** Quantification of n = 6; *p < 0.05, **p < 0.01 by one-way ANOVA with Tukey’s test.

Given that VRAC inhibitors and LRRC8A gene disruption reduce ischemic neuronal injury (Balkaya et al., 2023; Feng et al., 2004; Kimelberg et al., 2000; Kimelberg et al., 2003; Yang et al., 2019; Zhang et al., 2008; Zhou et al., 2020), we also investigated whether selective neuronal LRRC8A gene disruption blocks neuronal oxidative stress during cortical ischemia-reperfusion. Ischemia-reperfusion caused by transient middle cerebral artery occlusion (tMCAo) induced a robust increase in neuronal DHE oxidation of neurons in Wt mice, as previously reported (Kim et al., 2002) (Fig. 7), whereas this increase was blocked in the LRRC8A gene-disrupted neurons. Parallel findings were observed in using γH2Ax foci formation as a marker of intracellular oxidative stress (Fig. 7).

**Figure 7.**
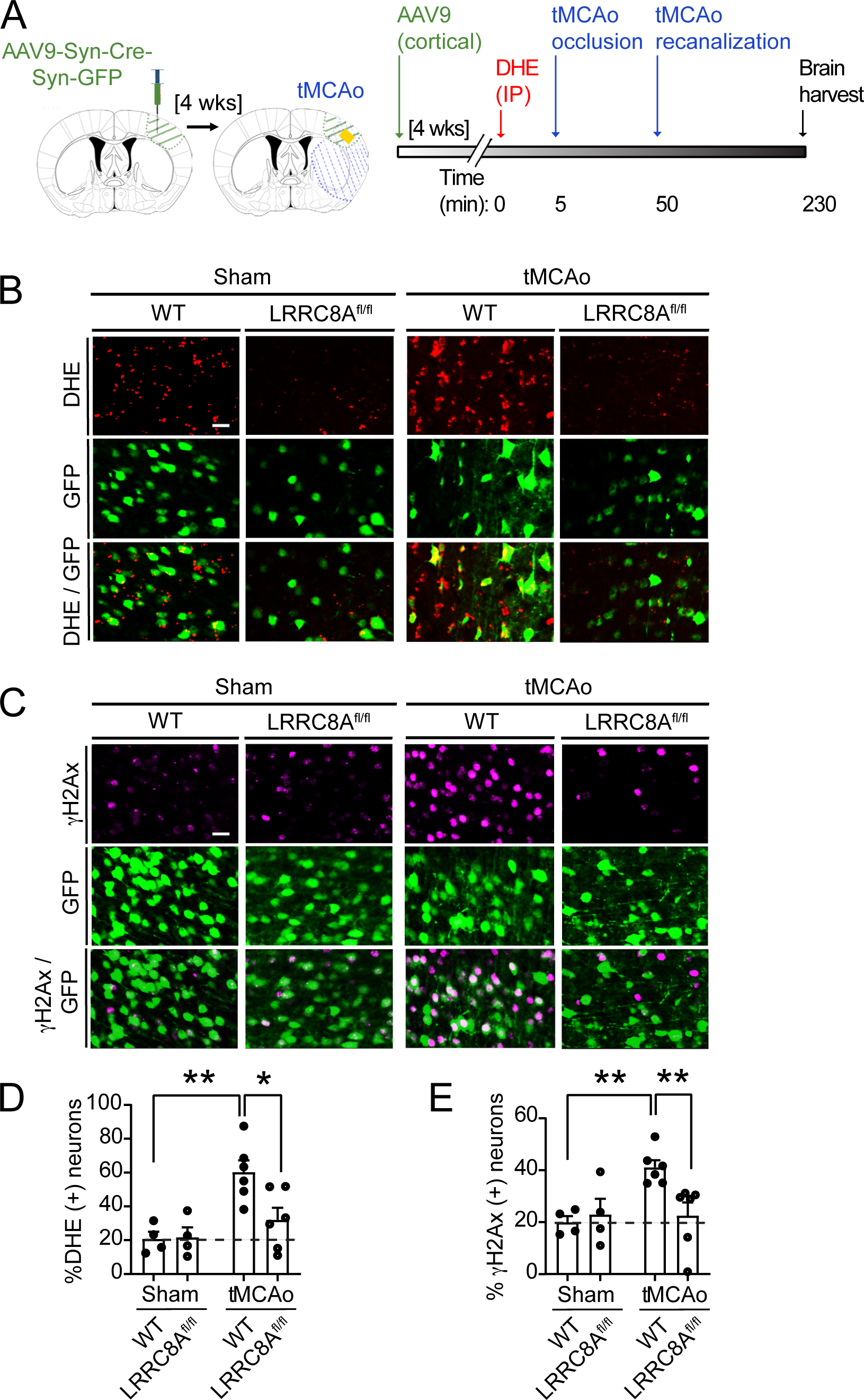
LRRC8A deletion prevents oxidative stress after transient MCA occlusion. **A.** Time course of AAV9, DHE, and NMDA injections. Red rectangle depicts the photographed region. **B,C.** Representative images of oxidized DHE (red) and yH2Ax foci (magenta) in AAV9-Syn-GFP-Syn-cre transduced neurons following tMCA occlusion or sham surgeries. Scale bar = 20 µm. **D,E.** Quantification of oxidized DHE and γH2Ax in the GFP(+) neurons. n = 6; *p < 0.05, **p < 0.01 by one-way ANOVA with Tukey’s test.

## DISCUSSION

Our findings show that extracellular superoxide crosses neuronal plasma membranes without prior conversion to H_2_O_2_; instead, it passes through LRRC8A-containing volume-regulated anion channels (VRACs). Using neuronal cultures we showed that extracellular superoxide generated by either KO_2_^-^ addition to the medium or activation of neuronal NMDA receptors led to elevated intracellular superoxide levels and concomitant DNA damage. The superoxide influx into neurons was blocked by superoxide dismutase, which converts superoxide to H_2_O_2_; by anion channel blockers including the putative VRAC-specific inhibitor DCPIB; and by Cre-lox disruption of the essential VRAC subunit, LRRC8A. Parallel results were observed in mouse brain, as increased neuronal superoxide levels and DNA damage were induced by NMDA injections or transient ischemia, and these were likewise blocked by both DCPIB and LRRC8A gene disruption.

VRACs are permeable to many anions, including chloride, taurine, glutamate, and others (Ghouli et al., 2022) but have not previously been identified as being permeable to superoxide. Superoxide anions are small relative to the estimated VRAC pore size of 1.2 - 1.4 nm in native lipid environments (Ghouli et al., 2022; Osei-Owusu et al., 2018), and given the broad range of anions gated by VRACs it is not surprising that superoxide is also conducted.

Consistent with the present findings, published brain atlas data show LRRC8A to be widely expressed in brain neurons (Yao et al., 2021). LRRC8 subunits B-D are also widely though variably expressed in neurons, whereas the LRRC8E subunit is not. The factors regulating VRAC opening varies with subunit composition of the channels as well as cell-type (Mongin, 2016 Planells-Cases et al., 2015; Sorensen et al., 2016; Gradogna et al., 2017). The factors that promote VDAC opening in neurons have not been well characterized, but in non-neuronal cell types these include ATP, G protein-coupled receptor (GPCR) stimulation, and redox state (Ghouli et al., 2022; Akita and Okada, 2011; Osei-Owusu et al., 2018). As VRACs are redox sensitive (Friard et al., 2021) it is possible that extracellular superoxide can activate as well as traverse VDACs VRACs can be inhibited by several compounds, including the anion channel blockers DIDS and DCPIB used here. DIDS is a non-specific chloride channel inhibitor, whereas DCPIB is considered a potent and selective inhibitor of VRACs (Gupta et al., 1991). DCPIB has also been reported to have off-target activity at concentrations used to inhibit VRACs (Afzal et al., 2019), but results obtained with DCPIB in the current study were very similar to those obtained by DIDS and genetic knockdown of LRRC8A.

We used DHE oxidation and γH2Ax formation as measures of superoxide entry into neurons and cellular oxidative stress, respectively. Dihydroethidium (DHE) is oxidized to fluorescent ethidium species that subsequently bind tightly to DNA and RNA and thus give an integrated measure over time of intracellular oxidant levels. γH2Ax is a phosphorylated form of the H2Ax histone, which accumulates at sites of double-stranded DNA breaks (Mah et al., 2010) and thereby provides a measure of intracellular oxidative stress. However, DHE oxidation and γH2Ax formation can also be induced by reactive oxygen species formed downstream from superoxide, such as hydroxyl radical, and hydrogen peroxide (H_2_O_2_). Since superoxide can form H_2_O_2_, and H_2_O_2_ can cross cell membranes, it was necessary to exclude the possibility that the extracellular superoxide was entering neurons in the form of H_2_O_2_. This was done by confirming that SOD negated the effects of extracellular superoxide, whereas catalase (which consumes H_2_O_2_) had minimal effect. Moreover, the oxidative signals induced by KO_2_ ^-^ were blocked by the anion channel inhibitors, whereas those induced by H_2_O_2_ were not.

KO_2_ ^-^ provides a useful way of exposing cells to controlled amounts of authentic superoxide, but under physiological conditions extracellular superoxide is produced primarily by NOX2. In neurons, NMDA receptor activation leads to extracellular superoxide production by NOX2 (Brennan-Minnella et al., 2013), and we thus used NMDA to stimulate this endogenous source of superoxide production. Our results showed that the effects of NMDA, like KO_2_ ^-^, were blocked by SOD, by VRAC blockade, and by LRRC8A downregulation. These findings demonstrate the physiological relevance of superoxide entry via VRACs. They also support prior observations indicating that that the oxidative stress induced by NMDA receptor activation is mediated primarily by release of superoxide into the extracellular space, rather than from intracellular mitochondria (Brennan-Minnella et al., 2013; Brennan et al., 2009; Clausen et al., 2013; Kishida et al., 2005; Reyes et al., 2012).

Glutamate excitotoxicity is a contributory factor in many neurological disorders, including ischemic stroke, with oxidative stress and DNA damage being proximate lethal events (Alano et al., 2010; Andrabi et al., 2011; Choi, 2020; Wang and Swanson, 2020; Wang et al., 2016). Our studies here using NMDA - induced excitotoxicity both in culture and in vivo showed that the resulting neuronal oxidative stress was blocked with either DCPIB or genetic LRRC8A downregulation, thus demonstrating a requirement for VRACs in the excitotoxic injury cascade. It was previously proposed that the protective effects of VRAC inhibitors and LRRC8A down-regulation in rodent models of stroke are attributable to reduced VRAC-mediated glutamate efflux from neighboring astrocytes (Feng et al., 2004; Kimelberg et al., 2000; Yang et al., 2019; Zhang et al., 2008; Zhou et al., 2020). Our finding that neuron-selective LRRC8A deletion blocks neuronal oxidative stress following ischemia or NMDA exposure does not directly contradict this idea, but a recent study showed a poor correlation between VRAC-mediated glutamate release and protection following MCAo in rodents (Balkaya et al., 2023). That study also showed that the deletion of VRACs solely in astrocytes was not sufficient to protect against ischemic injury, whereas deletion of VRACs in both neurons and astrocytes was protective (Balkaya et al., 2023).

Taken together, our results identify an essential role for VRACs in permitting the entry of superoxide into neurons and thus in excitotoxic injury. Accordingly, these results also suggest that determinants of neuronal VRAC localization and functional status likewise influence superoxide signaling and injury. As VRACs are increasingly recognized in diverse signaling pathways (Strange et al., 2019), a role for VRAC superoxide conductance in non-neuronal tissues is also likely.

## ACKNOWLEDGMENTS

This work was supported by grants from the NIH (NS081149; R.A.S.) and the Dept. of Veterans Affairs (BX004884; R.A.S.).

## AUTHOR CONTRIBUTIONS

Conceptualization, R.A.S.; Data collection, G.U, S-J.W. K.H., N.M, P.B, A.M, K.E; Writing – Original Draft, K.H., G.U., SJ-W; Writing – Review & Editing – R.A.S., K.H. Funding Acquisition, R.A.S.; Resources, R.S.; Supervision, R.A.S. and SJ-W.

## Supplemental Figure Legends

**Supplementary Figure 1. Catalase attenuates oxidative stress induced by H_2_0_2_, but not KO_2_^-^ in cultured neurons. A.** Oxidized DHE (red) in primary cortical neurons following incubation with KO_2_^-^ (40 μM) or H_2_O_2_ (100 μM). Cell nuclei are labeled with DAPI (blue). Where indicated, cultures were co-incubated with catalase (100 U / ml). **B.** γH2AX foci formation (green) in neurons treated as in A. **C,D.** Quantification of oxidized DHE and γH2AX. **p < 0.01 by Student’s t-test (n = 3).

**Supplementary Figure 2. Anion channel blockers do not prevent H_2_0_2_-induced oxidative stress in cultured neurons. A.** Oxidized DHE (red) in primary cortical neurons following incubation with H_2_O_2_ (100 μM). Cell nuclei are labeled with DAPI (blue). Where indicated, cultures were co-incubated with DCPIB (10 μM), or DIDS (100 μM). **B**. Foci of γH2AX formation (green) in neurons treated as in (A). Inset shows high magnification view of nuclei with γH2AX puncta. **C,D.** Quantification of oxidized DHE and γH2AX signals. **p < 0.01 versus H_2_O_2_ alone by Dunnett’s test (n = 3).

**Supplementary Figure 3. D.I.V. 7 primary cortical neurons do not exhibit oxidative stress in response to glutamate. A)** Oxidized DHE (red) in primary cortical neurons following incubation with glutamate (100 μM). H_2_O_2_ (100 μM) was used as a positive control. Cell nuclei are labeled with DAPI (blue). **B**. Foci of γH2AX formation (green) in neurons treated as in A. **C,D.** Quantification of oxidized DHE and γH2AX signals. **p < 0.01 versus H_2_O_2_ by Dunnett’s test (*n* = 3).

**Supplementary Figure 4. NMDA-induced oxidative stress is prevented by an inhibitor of NADPH oxidase-2**. Oxidized DHE (red) in primary cortical neurons following incubation with NMDA (100 μM). Where indicated, cultures were co-incubated with GSK2795039 (200 μM). Cell nuclei are labeled with DAPI (blue). **B**. Quantification of oxidized DHE. **p < 0.01 versus NMDA by Dunnett’s test (*n* = 3).

**Supplementary Figure 5. CLC-3 is does not permit superoxide entry in neurons.** Neurons prepared from mice lacking the ClC-3 chloride channel (*CLCn3*^-/-^ mice) were compared to wild-type neurons prepared in parallel. KO_2_^-^ and NMDA exposures were performed on D.I.V. 12. KO_2_^-^ and NMDA produced large elevations in DHE signal in both neuron types. There was no significant difference between the Wt and *ClCn3*^-/-^ neurons as assessed by ANOVA with Tukey’s test. Data are expressed relative to the wild-type neurons under the control condition in each experiment (n = 4).

