## Supplementary figures and images for "Superoxide enters neurons via LRRC8A – containing volume-regulated anion channels"

### Supplementary figure 1

Supplementary Figure 1

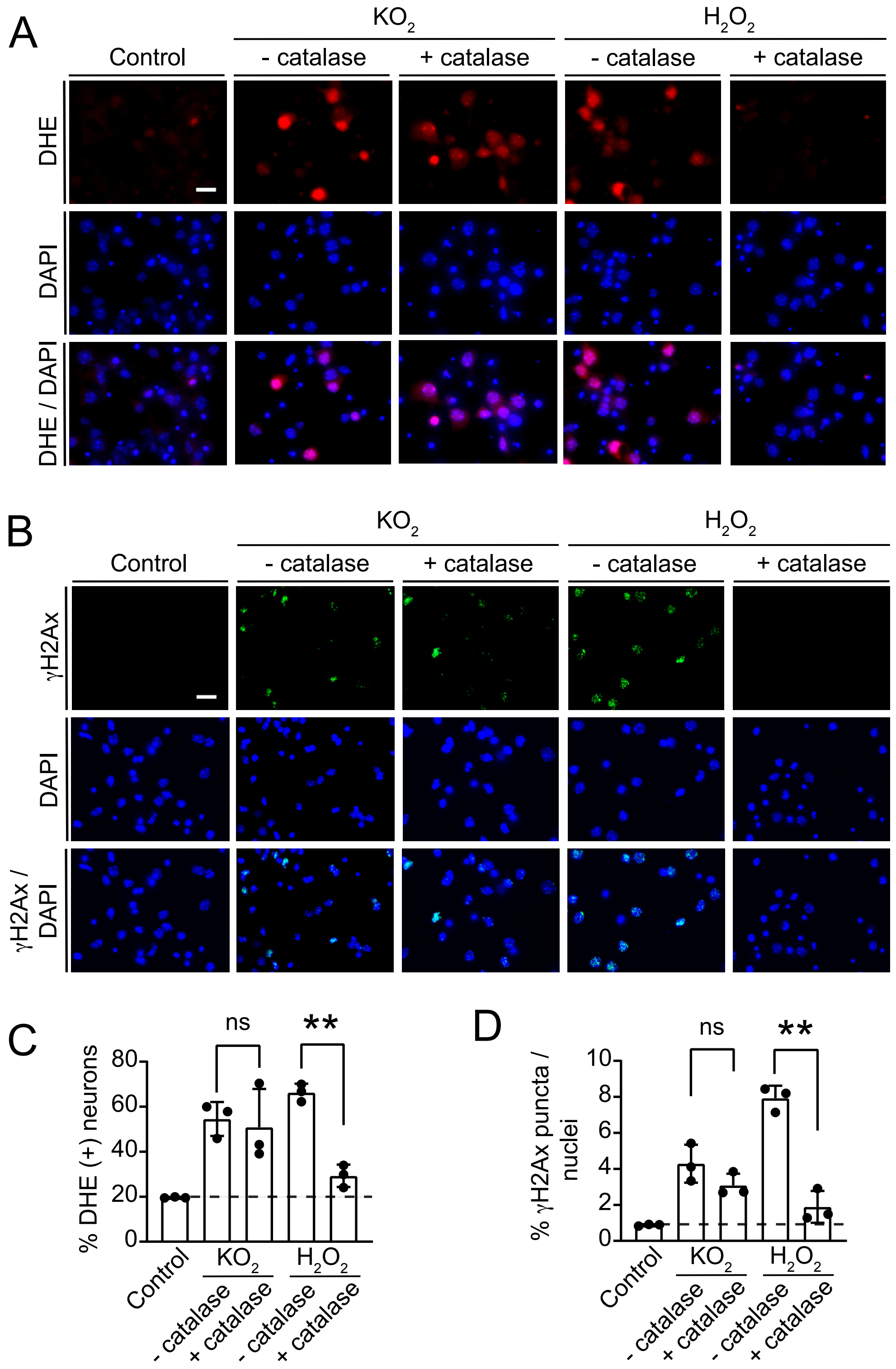

### Supplementary figure 2

Supplementary Figure 2

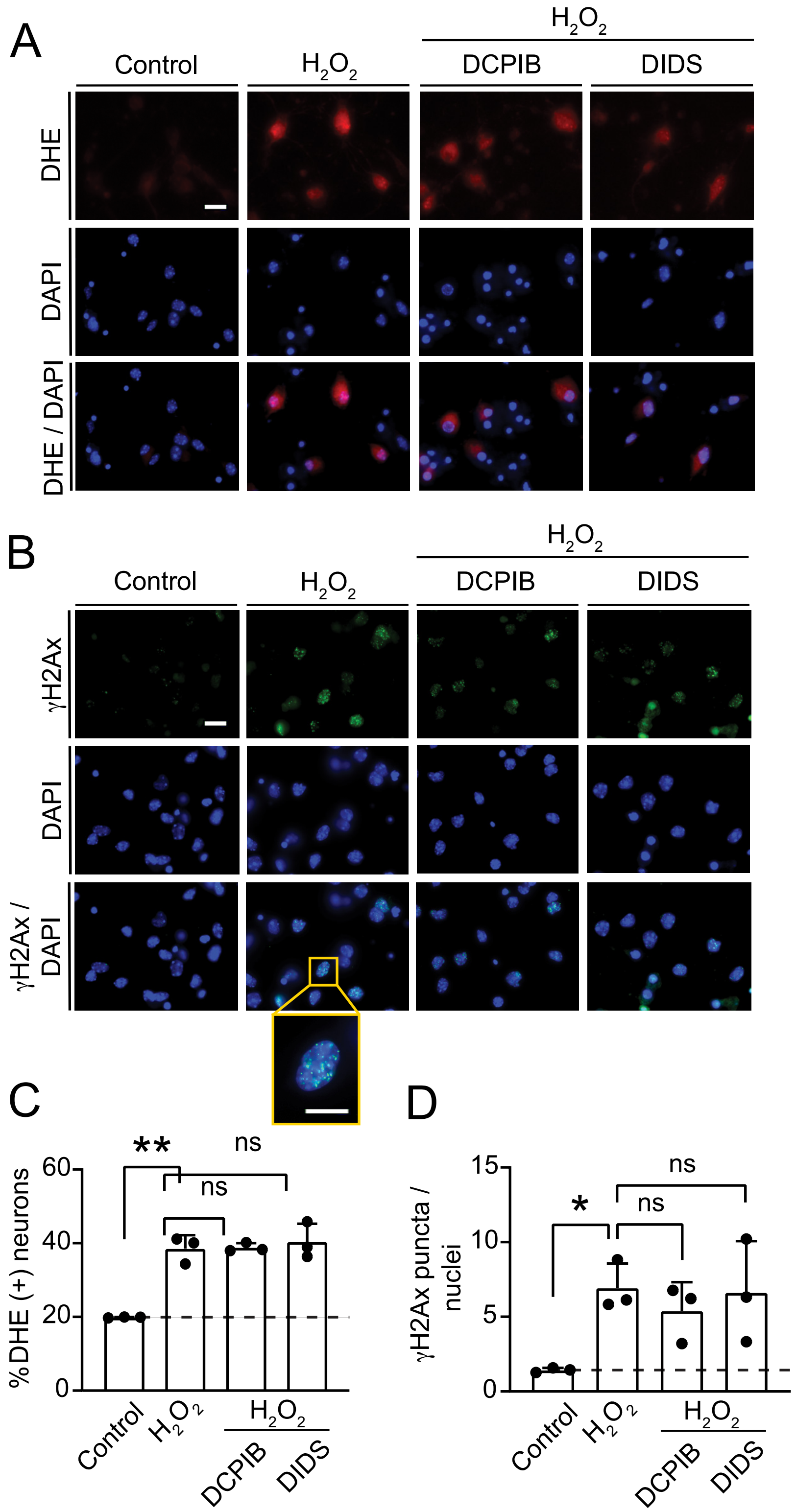

### Supplementary figure 3

Supplementary Figure 3

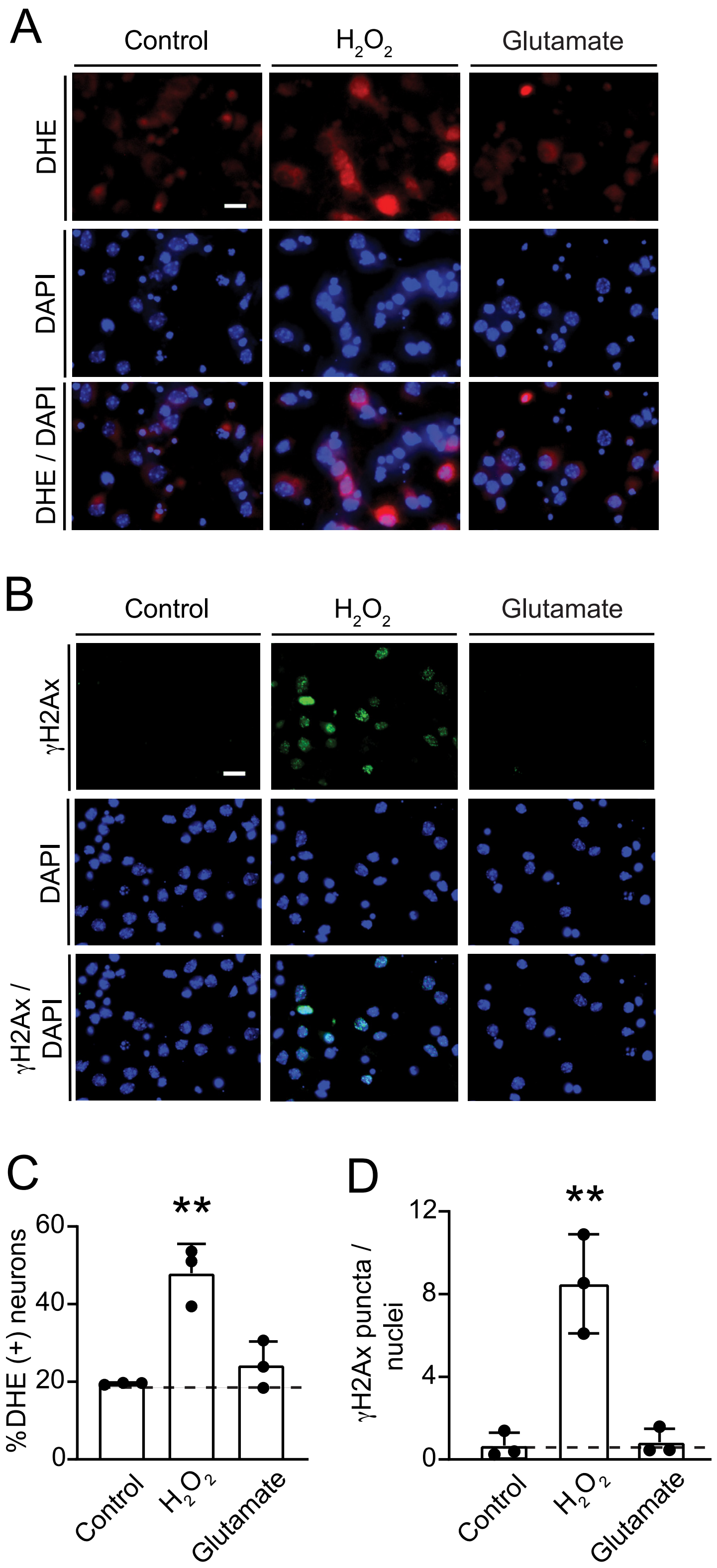

### Supplementary figure 4

Supplementary Figure 4

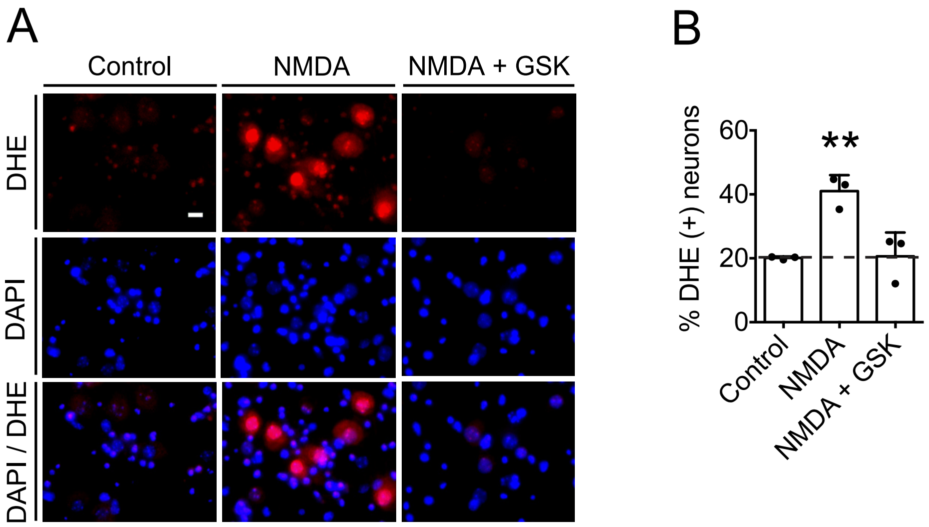

### Supplementary figure 5

Supplementary Figure 5

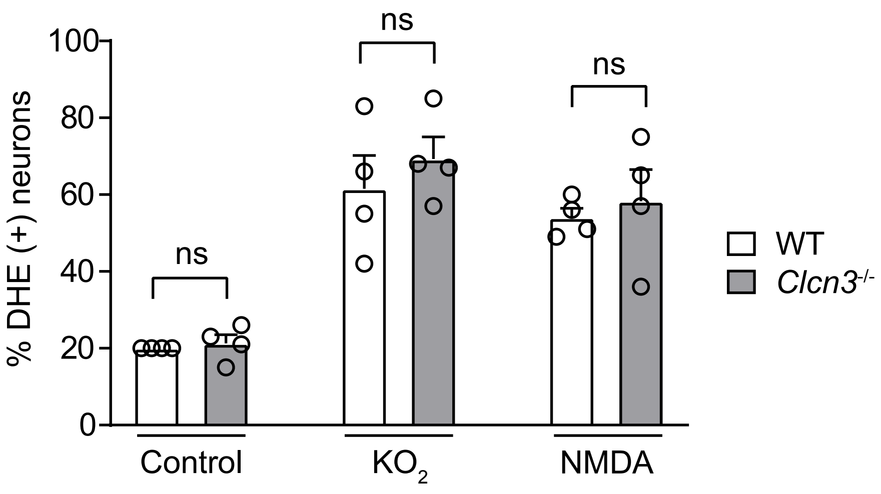
